# Adolescent stress impairs parvalbumin interneurons and their associated perineuronal nets: protective effects of microglia-modulating minocycline treatment

**DOI:** 10.1101/2025.09.03.674067

**Authors:** Victor Hadera, Bianca Caroline Bobotis, Antonia Landwehr, Francisco S. Guimarães, Marie-Ève Tremblay, Felipe V. Gomes

## Abstract

Adolescence is a critical period of brain maturation during which exposure to stress can lead to long-lasting behavioral and neurobiological alterations linked to increased vulnerability to psychiatric disorders. Here, we investigated whether minocycline, a tetracycline antibiotic that modulates microglial activity, could prevent or attenuate the long-term effects of adolescent stress on behavior, parvalbumin (PV)-expressing (+) interneurons (PVIs), perineuronal nets (PNNs), and microglia in adulthood. Male mice were exposed to a 10-day footshock stress protocol during adolescence (postnatal days 31–40) and treated with minocycline (30 mg/kg; i.p.) either during or after stress exposure. Behavioral assessments in adulthood revealed that adolescent stress impaired sociability, social memory, and object recognition memory, which were attenuated by minocycline treatment during or after adolescent stress exposure. Stress also reduced the number of PV+, PNN+, and PV+/PNN+ cells in the prefrontal cortex (PFC) and ventral hippocampus (vHip). These effects were prevented by minocycline administration at both time points. No significant long-lasting changes were observed in microglial number, density, or spatial distribution in either region. However, minocycline treatment modulated microglial morphology in a region- and timing-dependent manner, with increased microglial area observed in the PFC and subtle alterations in circularity in the vHip. These findings suggest that adolescent stress induces enduring impairments in PVIs and behavior, possibly through transient microglial intervention and PNN degradation. Minocycline treatment during or after stress was effective in preventing these changes, supporting its potential as a therapeutic strategy to mitigate the long-term consequences of adolescent stress and to reduce vulnerability to stress-related psychiatric disorders.

## INTRODUCTION

The rise in mental disorders, such as anxiety and depression, represents a significant challenge for public health due to their complex and multifactorial etiology [1,2]. Exposure to adverse socio-environmental factors during critical periods of neurodevelopment, such as adolescence, increases the risk of developing psychiatric disorders later in life [3–6].

Adolescence is a period of intense neuroplasticity, during which traumatic experiences can impact critical brain regions involved in emotional and cognitive functions, including the prefrontal cortex (PFC) and hippocampus [7–9]. Animal studies have shown that rodents exposed to stressors during a period corresponding to adolescence in humans exhibit long-lasting deficits in sociability and cognitive function, along with morphological changes in the brain [10–15]. Among these changes, deficits in (PV)-expressing (+) GABAergic interneurons (PVIs) have been reported in both the PFC and ventral hippocampus (vHip) [10,13–15].

Due to their fast-spiking activity, PVIs are critical in regulating the excitatory-inhibitory balance [16,17]. During adolescence, PVIs are still undergoing maturation [18], which coincides with the full formation of a specialized extracellular matrix, the perineuronal nets (PNNs), that enwrap most PVIs [19–21]. In addition to stabilizing synaptic inputs onto PVIs [22], PNNs protect them from damage induced by redox dysregulation [23,24] and exposure to psychological stress [14]. It has been proposed that, during adolescence, when PVIs are not fully mature and protected by PNNs, they may be particularly vulnerable to the deleterious effects of stress [14,25].

There is also evidence that stress modulates microglia, the brain’s resident immune cells, which play key roles in synaptic pruning and stress response [26]. Microglial dysfunction has also been associated with PNN degradation [27–30], which may contribute to a functional loss of PVIs and, consequently, disruption of the excitatory-inhibitory balance.

Minocycline, a tetracycline antibiotic, has shown potential in modulating microglial activity [31] and attenuating the effects of stress by reducing the release of pro-inflammatory cytokines and modulating inflammatory signaling pathways [32]. Clinical and preclinical studies suggest that minocycline may have benefits in the treatment of various psychiatric disorders such as anxiety, depression, and schizophrenia. However, its impact on the effects of stress during adolescence remains underexplored.

Here, we investigate in mice whether treatment with minocycline during or after exposure to stress during adolescence mitigates its long-lasting effects on behavior, PVIs and their associated PNNs, as well as morphological changes in microglia (Iba1+ cells) within the PFC and vHip into adulthood.

## METHODS

### Animals

Male C57BL/6J mice (3 weeks old) were obtained from the Central Animal Facility of the University of São Paulo, Ribeirão Preto campus. The animals were housed in groups of 3-4 per cage in a temperature-controlled room (24 ± 1°C) with a 12-hour light/dark cycle (lights on at 6:30 AM), with food and water *ad libitum*. All procedures were approved by the Ethics Committee of the Ribeirão Preto Medical School (#49/2021), which follows Brazilian and international guidelines for animal care and use. Of note, only male mice were used, as previous findings showed that female adolescent mice were resistant to long-lasting behavioral deficits following exposure to the same stress protocol [15].

### Adolescent stress protocol

Animals were subjected to a stress protocol consisting of daily 15-minute sessions of footshock exposure (15 shocks, 0.75 mA, 2 seconds) for 9 consecutive days, from postnatal day (PND) 31 to 40 [15]. Naïve animals were kept undisturbed in the animal room. Behavioral tests to assess social interaction, cognitive function, as well as the morphological analyses, were performed in adulthood (PND62-64).

### Minocycline treatment

Initially, we evaluated the effects of minocycline treatment during exposure to stress (n=8/group). Animals were subjected to the adolescent stress protocol between PND 31–40, and one hour after each stress session, they received an intraperitoneal injection (i.p.) of either minocycline (30 mg/kg; Sigma) or saline (vehicle; 10 mL/kg). To evaluate the effects of minocycline after adolescent stress, in a second cohort, the administration of minocycline (30 mg/kg, i.p.) or saline was initiated one day after the end of the stress protocol and continued for 10 days (PND 41–50; n=8/group). This treatment dosage has been previously used in other studies in mice from PND 30 to 40 [33].

### Behavioral tests

#### Social interaction and social discrimination in the three-chamber test

On PND62, animals were tested for sociability and social discrimination. The apparatus consisted of an acrylic box with three interconnected compartments (20 cm x 40 cm each). Test animals were first habituated to the apparatus for 5 min, with two empty wire cages (i.e., no “interaction” animal) placed in the lateral compartments. To evaluate social interaction, an unfamiliar mouse of the same strain, sex, and age was placed inside a cage in one of the side compartments, while the other cage remained empty. The test animal, placed in the central compartment, was allowed to explore the apparatus for 10 min. Next, to assess social discrimination, the test mouse was confined in the central compartment for 1 min. The previously introduced unfamiliar mouse was now designated as the familiar social stimulus, and a novel, unfamiliar mouse was placed in the opposite cage. The test animal was then allowed to explore the apparatus for an additional 5 min. Behaviors such as sniffing, touching, and climbing on the cages were recorded. Results were reported as total interaction time (in seconds), as well as the Social Interaction (SI) and Social Discrimination (SD) indices, calculated using the following formulas:

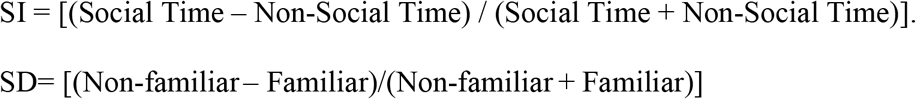

#### Novel Object Recognition (NOR) Test

On PND62-63, novel object discrimination was assessed in the NOR test, which was conducted in a circular acrylic arena (40 cm in diameter and 40 cm in height). One day before testing, on PND62, the animals were habituated to the arena for 15 min. On the test day (PND63), during the first session (acquisition phase, T1), animals were placed in the arena containing two identical objects and allowed to explore for 10 min. One hour later, during the second session (retention phase, T2), one of the objects from the first session was replaced with a novel object (NO), placed in the same location as in T1. Animals were then allowed to explore the arena for 5 min. Object exploration was defined as the animal orienting its face toward the object at an approximate distance of at least 2 cm while observing, sniffing, or touching it. Recognition memory was assessed using the discrimination index (DI), calculated as follows: DI = [(novel object – familiar object) / (novel object + familiar object)].

### Immunofluorescence for PV, PNNs, and Iba1 in the PFC and vHip

On PND64, one day after behavioral testing, animals were anesthetized with chloral hydrate (400 mg/kg; i.p.) and transcardially perfused with 0.01M phosphate-buffered saline (PBS), followed by 4% paraformaldehyde (PFA). Brains were removed, post-fixed in 4% PFA for 24 hours, and transferred to a 30% sucrose solution for 30 hours. For each animal, 5 to 6 coronal sections (30 µm thick), spanning the rostrocaudal axis, containing the PFC (+1.98 – +1.70 mm from bregma) and vHip (−2.92 – −3.80 mm from bregma), were collected and processed using the free-floating method to assess the expression of PV, PNN, and Iba1 via immunofluorescence. Sections were incubated for 18 hours at 4°C with a solution containing 1% fetal bovine serum, mouse anti-PV antibody (1:1000, Sigma, #P-3088), biotinylated Wisteria floribunda agglutinin (WFA, 1:500, VectorLaboratories, #FL-1351-2), and rabbit anti-Iba1 antibody (1:1000, Wako, #019-19741). Subsequently, sections were incubated in a secondary antibody solution containing 1% fetal bovine serum, goat anti-mouse IgG conjugated to AlexaFluor 647 (1:1000, Thermo Scientific, #A-21235), goat anti-rabbit IgG conjugated to AlexaFluor 488 (1:1000, Thermo Scientific, #A-11008), and streptavidin conjugated to AlexaFluor 594 (1:1000, Thermo Scientific, #S32356). Sections were also counterstained with DAPI (1:4000, Abcam, #ab104139).

#### Quantification of PV+, PNN+, PV+/PNN+, and Iba1+ cells

For imaging, the focal plane was adjusted using the PV+ cell channel, and all subsequent images were captured using identical exposure settings across groups. Images were acquired using a Leica LAS X SP8 confocal microscope with 20× (numerical aperture = 1.3) and 40× objectives (numerical aperture =1.4). Z-stacks were collected with an interval of 26 µm for 20X objective and 19 µm for the 40X objective between each section, averaging 43 stacks/image for the 20X objective and 20 stacks/image for the 40X objective. Cell counting was performed on images acquired at 20X magnification using the Cell Counter Plugin in ImageJ software (NIH, v.1.8.0_322). PV+, PNN+, PV+/PNN+, and Iba1+ cells located within the boundaries of the prelimbic and infralimbic portion of the PFC and the subicular region of the vHip were manually counted, regardless of fluorescence intensity. In addition to quantifying Iba1+ microglia, we evaluated the spatial distribution of these cells using the nearest neighbor distance (NND; measured in μm), which indicates whether cells are uniformly distributed, clustered, or dispersed. We also calculated the spacing index (reported in arbitrary units, AU), defined as NND^2^ multiplied by Iba1+ cell density. This parameter combines the proximity and the total number of cells, providing further insight into whether Iba1+ microglia are more dispersed or densely packed within the analyzed regions [34,35].

#### Automatic morphological analysis of individual Iba1+ cells

Microglia morphology analyses were performed on images acquired at 40X magnification using the CellProfiler software (Broad Institute; MIT/Harvard, v.4.2.6) [36,37]. The images were blinded to the experimental groups and 336 images of Iba1+ cells from the vHip and PFC from 8 mice/group (a total of 64 mice) were run in the pipeline. The pipeline consisted of microglial detection by determining a pixel size range (43-350) and threshold strategies (Robust Background) allowing the software to accurately recognize its soma and processes. After the detection, morphological modules, such as ‘Morph’, ‘MeasureObjectSkeleton’ and ‘MeasureObjectSizeShape’ were added to provide in depth measurements of the previously detected microglial objects. In addition to the number of trunks, the outputs generated area, circularity, and skeleton length measured in pixels/AU [38].

### Statistical analyses

Statistical analyses were conducted using SPSS (IBM SPSS Statistics, v. 22.0) and GraphPad Prism (v. 10). Data were analyzed using a two-way analysis of variance (ANOVA), with condition (naïve or stressed) and treatment (vehicle or minocycline) as main factors, followed by Tukey’s *post-hoc* test. Normality was tested using the Shapiro–Wilk test. All parameters passed normality except the Iba1+ microglia morphology results (including microglial cell area, circularity, number of trunks, and total skeleton length). These data were then analyzed using the non-parametric Kruskal-Wallis test, followed by Dunn’s *post-hoc* test. Statistical significance was set at p<0.05. No outliers were excluded from the analyses.

## RESULTS

### Treatment with minocycline during and after the end of adolescent stress attenuates behavioral deficits

In adulthood, animals exposed to adolescent stress exhibited decreased social interaction, social discrimination, and object recognition memory. All these behavioral deficits were mitigated by minocycline treatment initiated either during or after the stress protocol (Figure 1).

**Figure 1.**
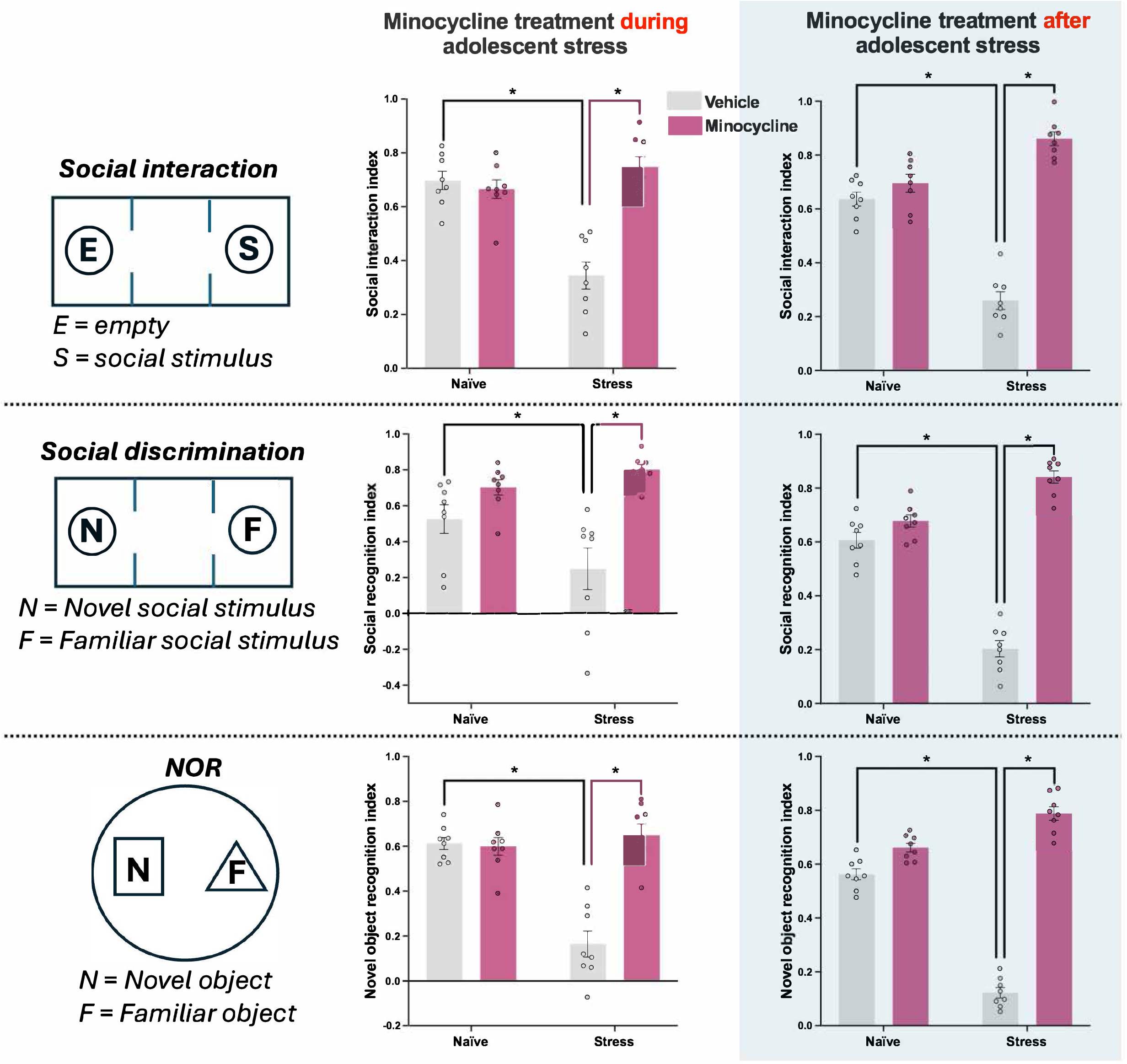
Minocycline treatment during or after adolescent stress prevents deficits in social interaction, social discrimination and novel object discrimination in adult mice. Adult mice (PD 62-63) previously exposed to adolescent stress (PD 31-40) showed reduced social interaction and social discrimination in the three-chamber test, along with impairment in object discrimination in the novel object recognition (NOR) test. Minocycline treatment (30 mg/kg) administered either during (PD 31-40) or after (PD 41-50) the stress protocol attenuated these deficits (n=8/group). Data are shown as mean ± SEM. *p<0.05, two-way ANOVA followed by Tukey’s *post-hoc* test.

When minocycline treatment was administered during adolescent stress, two-way ANOVA revealed *condition* x *treatment* interactions for the social interaction [F(1,28)=29.82; p<0.05], social discrimination, [F(1,28)=6.32; p<0.05], and novel object discrimination [F(1,28)=30.22; p<0.05] indices. Tukey’s *post-hoc* test revealed that stressed animals presented lower social interaction, social discrimination, and novel object discrimination (p<0.05 vs. naïve + vehicle), indicating, respectively, impaired sociability, social memory, and object recognition memory. These changes were prevented by minocycline treatment during stress (p<0.05 vs. stress + vehicle; Figure 1).

Similarly, when minocycline treatment was administered after adolescent stress, our analyses indicated once more *condition* x *treatment* interactions for the social interaction [F(1,28)=85.74; p<0.05], social discrimination, [F(1,28)=113.2; p<0.05], and novel object discrimination [F(1,28)=190.5; p<0.05] indices. After Tukey’s *post-hoc* test, a decrease in social interaction, social discrimination, and novel object discrimination caused by adolescent stress (p<0.05 vs. naïve + vehicle), which were mitigated by minocycline treatment post-stress (p<0.05 vs. stress + vehicle; Figure 1).

### Adolescent stress decreases the number of PV+ and PNN+ cells in the PFC and vHip, which is attenuated by minocycline treatment during or after stress

Adult mice exposed to stress during adolescence presented decreased numbers of PV+, PNN+, and PV+/PNN+ cells in both the PFC and vHip. These changes were mitigated by the treatment with minocycline administered either during or post-stress (Figure 2).

**Figure 2.**
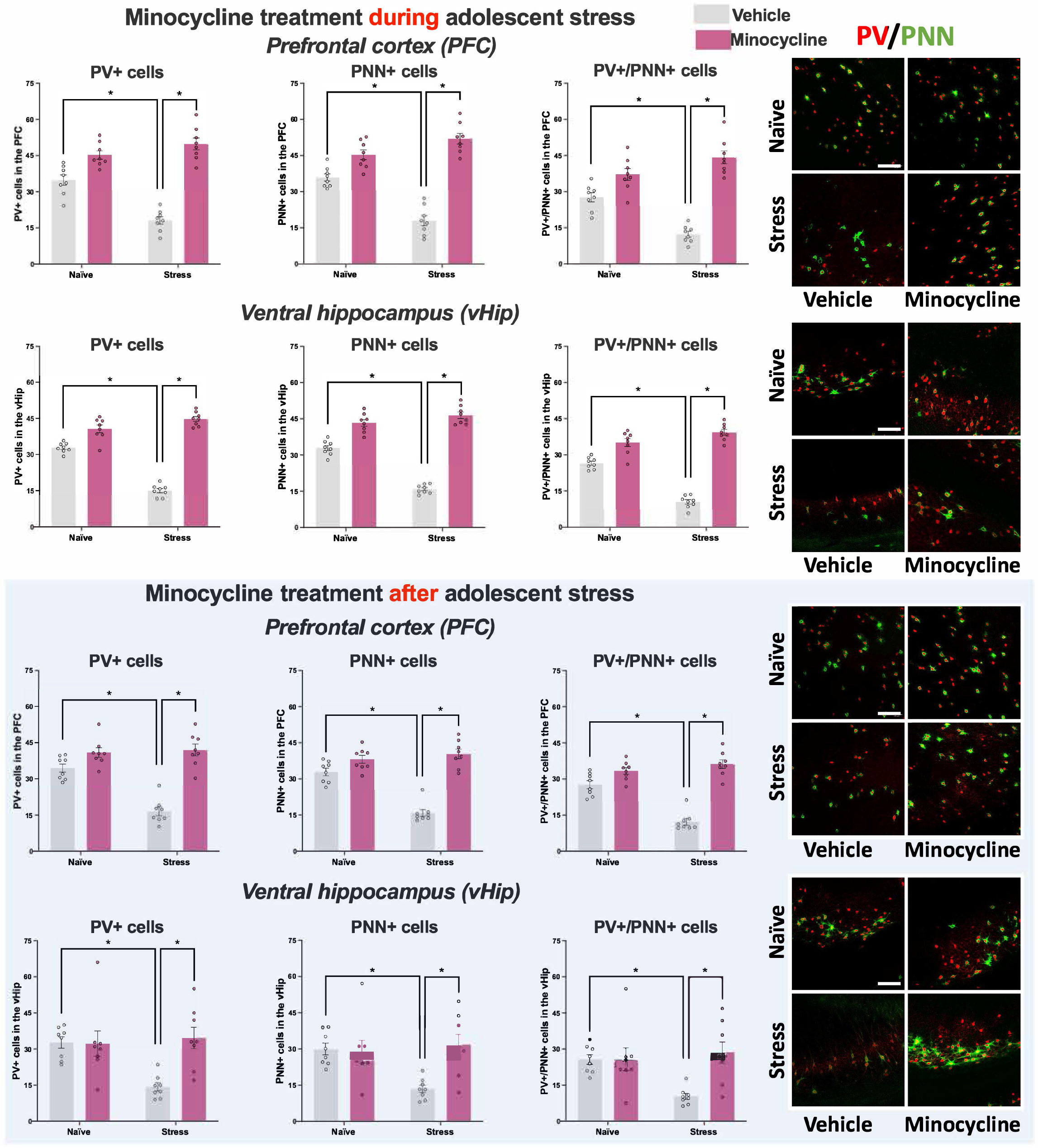
Minocycline treatment during or after adolescent stress prevents decreases in the number of PV+, PNN+, and PV+/PNN+ cells in the prefrontal cortex (PFC) and ventral hippocampus (vHip) in adult mice. Exposure to adolescent stress (PD31-40) caused a reduction in the number of PV+ (red), PNN+ (green), PV+/PNN+ cells in the PFC and vHip during adulthood. Minocycline treatment (30 mg/kg), administered either during (PD31-40) or after (PD41-50) the adolescent stress protocol, mitigated these reductions (n=8/group). Data are shown as mean ± SEM. *p<0.05, two-way ANOVA followed by Tukey’s *post-hoc* test. Scale bar: 100 μm.

#### PFC

In the PFC, when we evaluated the effects of minocycline treatment administered during the adolescent stress protocol, two-way ANOVA revealed *condition* x *treatment* interactions for the number of PV+ [F(1,28)=28.19; p<0.05], PNN+ [F(1,28)=39.56; p<0.05], and PV+/PNN+ cells [F(1,28)=27.65; p<0.05]. Tukey’s *post-hoc* test indicated reductions in the number of PV+, PNN+, and PV+/PNN+ cells in stressed animals (p<0.05 vs. naïve + vehicle), which were prevented by minocycline treatment during stress (p<0.05 vs. stress + vehicle; Figure 2).

In the experiment evaluating the effects of minocycline after adolescent stress, in the PFC, our analyses also indicated *condition* x *treatment* interactions for the number of PV+ [F(1,28)=22.23; p<0.05], PNN+ [F(1,28)=33.58; p<0.05], and PV+/PNN+ cells [F(1,28)=33.96; p<0.05]. *Post-hoc* comparisons confirmed reductions in the number of PV+, PNN+, and PV+/PNN+ cells in stressed animals (p<0.05 vs. naïve + vehicle), which were also attenuated by minocycline treatment administered after stress (p<0.05 vs. stress + vehicle; Figure 2).

#### vHip

Similar to the PFC, adolescent stress reduced the number of PV+, PNN+, and PV+/PNN+ cells in the vHip. For minocycline treatment during adolescent stress, two-way ANOVA indicated *condition* x *treatment* interactions for the number of PV+ [F(1,28)=98.77; p<0.05], PNN+ [F(1,28)=77.45; p<0.05], and PV+/PNN+ cells [F(1,28)=72.35; p<0.05]. Tukey’s *post-hoc* test showed reductions in the number of PV+, PNN+, and PV+/PNN+ cells after adolescent stress (p<0.05 vs. naïve + vehicle), which were prevented by minocycline treatment during adolescent stress (p<0.05 vs. stress + vehicle; Figure 2).

For animals treated with minocycline after stress, two-way ANOVA also indicated *condition* x *treatment* interaction for PV+ [F(1,28)=7.73; p<0.05], PNN+ [F(1,28)=7.63; p<0.05], and PV+/PNN+ cells [F(1,28)=6.92; p<0.05]. Again, Tukey’s *post-hoc* test revealed that PV+, PNN+, and PV+/PNN+ cells diminished in the vHip after adolescent stress (p<0.05 vs. naïve + vehicle). These reductions were also attenuated by minocycline treatment post-stress (p<0.05 vs. stress + vehicle; Figure 2).

### Effects of adolescent stress and minocycline treatment during or after stress on the number and morphology of Iba1+ microglia in the PFC and vHip of adult animals

#### Iba1-positive cell number and distribution

Adolescent stress and minocycline treatment, regardless of timing, did not alter the number of Iba1+ cells in the PFC or vHip. Similarly, no significant differences affected microglia regarding cell density, NND, or the spacing index in either brain region (Supplementary figure 1).

#### Iba1+ cells morphology

For the experiment evaluating the effects of minocycline administered during adolescent stress, the analysis of Iba1+ microglial morphology in the PFC revealed changes in circularity (H=7.74; p<0.05), number of trunks (H=9.19; p<0.05), and total skeleton length (H=9.21; p<0.05), with no changes in cell area. Dunn’s *post-hoc* test indicated that PFC microglia from minocycline-treated stressed animals showed increased circularity and skeleton length, along with fewer trunks, compared to those from controls (p<0.05 vs. naïve + vehicle; Figure 3). In the vHip, changes were observed in circularity (H=10.37; p<0.05) and skeleton length (H=7.68; p<0.05), with no changes in cell area or number of trunks. Microglia from minocycline-treated stressed animals exhibited increased circularity (Dunn’s *post-hoc* test; p<0.05 vs. naïve + vehicle), while microglia from minocycline-treated naïve animals showed greater skeleton length (Dunn’s *post-hoc* test; p<0.05 vs. naïve + vehicle; Figure 3).

**Figure 3.**
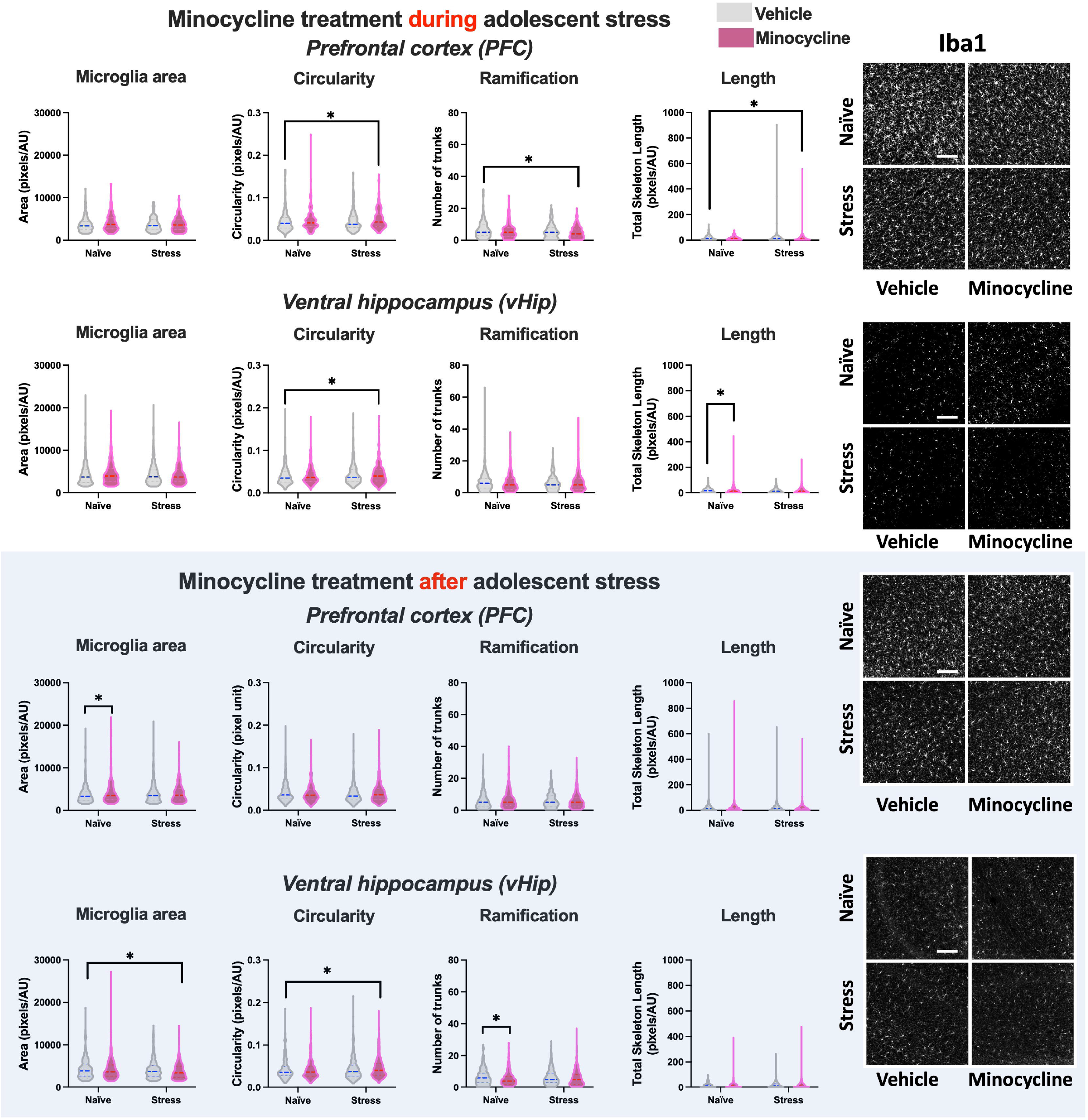
Effects of adolescent stress and minocycline treatment during or after stress on Iba1+ microglial morphology in the prefrontal cortex (PFC) and ventral hippocampus (vHip) of adult mice. For animals treated during stress, in the PFC, microglia from minocycline-treated stressed animals showed increased circularity and skeleton length, and fewer trunks than control animals (naïve + vehicle). In the vHip, circularity was increased in minocycline-treated stressed animals, while minocycline-treated naïve animals showed greater skeleton length (PFC – naïve + vehicle: 216 cells; naïve + minocycline: 201 cells; stress + vehicle: 170 cells; stress + minocycline: 203 cells; vHip – naïve + vehicle: 461 cells; naïve + minocycline: 408 cells; stress + vehicle: 442 cells; stress + minocycline: 436 cells. Values represent the total number of cells analyzed from 8 animals/group). For animals treated after stress, changes in microglial area were observed in the PFC, with increased area in minocycline-treated naïve animals. In the vHip, minocycline-treated stressed animals showed reduced area and increased circularity, and minocycline-treated naïve animals exhibited fewer trunks (PFC – naïve+vehicle: 468 cells; naïve + minocycline: 445 cells; stress + vehicle: 477 cells; stress + minocycline: 456 cells; vHip – naïve + vehicle: 312 cells; naïve + minocycline: 375 cells; stress + vehicle: 423 cells; stress + minocycline: 407 cells. Values represent the total number of cells analyzed from 8 animals/group). Data from individual cells are presented as violin plots and were analyzed using Kruskal–Wallis tests followed by Dunn’s *post-hoc* test. Units are reported as pixels/arbitrary units (AU). Scale bar: 100 μm.

In the experiment evaluating the effects of minocycline after adolescent stress, a change in microglial area was found in the PFC (H=6.96; p<0.05), with microglia from minocycline-treated naïve animals showing greater cell area (Dunn’s *post-hoc* test; p<0.05 vs. naïve + vehicle; Figure 3). However, no changes in circularity, number of trunks, and skeleton length were found. In the vHip, changes were found in cell area (H=9.53; p<0.05), circularity (H=9.63; p<0.05), and number of trunks (H=10.44; p<0.05), with no change in skeleton length. Dunn’s *post-hoc* test indicated that microglia from minocycline-treated stressed animals presented lower cell area along with greater circularity compared to those from controls (p<0.05 vs. naïve + vehicle; Figure 3). Additionally, microglia from minocycline-treated naïve animals presented fewer trunks (Dunn’s *post-hoc* test; p<0.05 vs. naïve + vehicle; Figure 3).

## DISCUSSION

Our results indicate that adolescent stress leads to long-lasting impairments in sociability, social memory, and object recognition memory in adult male mice, along with a reduction in PV+ cells, including those surrounded by PNNs, in the PFC and vHip. These findings are consistent with those of a previous study from our group [15]. However, in addition to the PFC, we now show that adolescent stress-induced deficits in PV+, PNNs+, and PV+/PNNs+ cells also extend to the vHip in mice. Minocycline treatment, whether administered during the 10 days of adolescent stress exposure (PND31-40) or in a 10-day regimen starting the day after stress cessation (PND41-50), attenuated both behavioral and cellular deficits in adulthood.

Consistent with previous studies [10,13–15,39], rodents exposed to adolescent stress presented impairments in social behavior and cognitive performance in adulthood, reinforcing that adolescence represents a critical period of vulnerability to environmental insults. The observed reductions in PV+ and PNN+ cells in the PFC and vHip align with studies reporting that stress applied during sensitive periods, such as childhood and adolescence, can impair the maturation and integrity of PVIs and their associated PNNs [10,13,15,40], which are essential for maintaining the excitatory-inhibitory balance and supporting cognitive and social functioning [41,42]. These data support the notion that stress-induced dysfunction in PVIs and PNNs may drive behavioral deficits relevant to psychiatric disorders, such as schizophrenia and depression [43–45]. However, the underlying mechanisms associated with adolescent stress-induced deficits in PVIs and PNNs remain unknown.

Given that stress can modulate microglia and disrupt synaptic plasticity [46,47], potentially contributing to PNN loss and an altered excitatory/inhibitory balance due to PVI dysfunction, we assessed whether adolescent stress induces long-lasting changes in the density and morphology of microglia in the PFC and vHip. In addition to the reduced presence of PNNs surrounding PVIs during adolescence, which has been proposed to contribute to the increased vulnerability of PVIs to stress, reactive microglia could further degrade PNNs [30], disrupt their developmental trajectory, and contribute to long-lasting behavioral changes.

Our findings showed that minocycline, which attenuates microglial reactivity [48], prevented adolescent stress-induced behavioral deficits and reductions in the number of PV+, PNN+, and PV+/PNN+ cells in both the PFC and vHip, regardless of whether it was administered during or after the stress period. This suggests that both prophylactic and early post-stress interventions with minocycline may be effective in mitigating long-term abnormalities caused by adolescent stress exposure. Supporting our results, studies using lipopolysaccharide (LPS)-induced microglial stimulation reported reduced PV expression in the PFC of rats, accompanied by cognitive impairments, both reversed by minocycline treatment [49]. In addition, microglial modulation by minocycline during chronic stress exposure of adult rats and mice has also been shown to improve stress-induced behavioral deficits [32,50].

Although stress can induce transient microglial changes in several brain regions associated with emotional regulation [46,51], we found no long-lasting alterations in Iba1+ cell number, density, spatial distribution, or gross morphology in the PFC and vHip of adult animals exposed to adolescent stress. However, subtle changes in microglial cell area and circularity were detected, depending on brain region and treatment timing. In the PFC, minocycline treatment during or after stress increased microglial cell area without altering other morphological parameters. This may reflect a shift toward a more surveillant or metabolically active microglial state, possibly related to increased synaptic support or clearance of stress-related debris [52,53]. In contrast, in the vHip, post-stress minocycline treatment reduced the Iba1+ microglial area, while stress exposure led to an increase in microglial circularity, an effect not reversed by minocycline. These modest alterations may reflect transient stress responses that occurred earlier and were no longer detectable in adulthood. However, even modest changes could have an impact on their function, which would warrant further investigation.

The absence of marked persistent microglial changes caused by adolescent stress aligns with previous reports showing that stress-induced microglial reactivity may normalize over time. For instance, although the number of Iba1+ cells increased 2 days after acute stress in adult mice, no changes were observed 4 days later [54]. Moreover, the microglial response to stress depends on factors such as type, duration, and timing of stress exposure [55,56]. While stress may modulate microglia, our findings suggest that adolescent stress does not lead to persistent major microglial changes in adulthood. However, we cannot rule out the possibility that adolescent stress transiently increases microglial reactivity, which could lead to functional changes, such as contributing to the degradation of PNNs and the subsequent dysfunction of PVIs. By dampening excessive microglial activation during or shortly after adolescent stress exposure, minocycline may prevent this early response, preserving PNN formation/integrity and PVIs, thereby mitigating behavioral deficits. This remains to be further investigated.

In conclusion, our results show that adolescent stress disrupts PVIs and their associated PNNs in the PFC and vHip and leads to behavioral impairments in adulthood. Minocycline treatment, both during and after stress, prevents these alterations, supporting its potential as a therapeutic strategy for preventing stress-related psychiatric disorders.

## ACKNOWLEDGMENTS

The authors thank Marco Antonio de Carvalho, Eleni Tamburus Gomes, and Eliane Aparecida Antunes Maciel for technical assistance. We also thank Adriano José Maia Chaves Filho for the help with the analysis of Iba1+ microglial counts and spatial distribution.

