## Supplementary figures for "Adolescent stress impairs parvalbumin interneurons and their associated perineuronal nets: protective effects of microglia-modulating minocycline treatment"

**Supplementary figure**

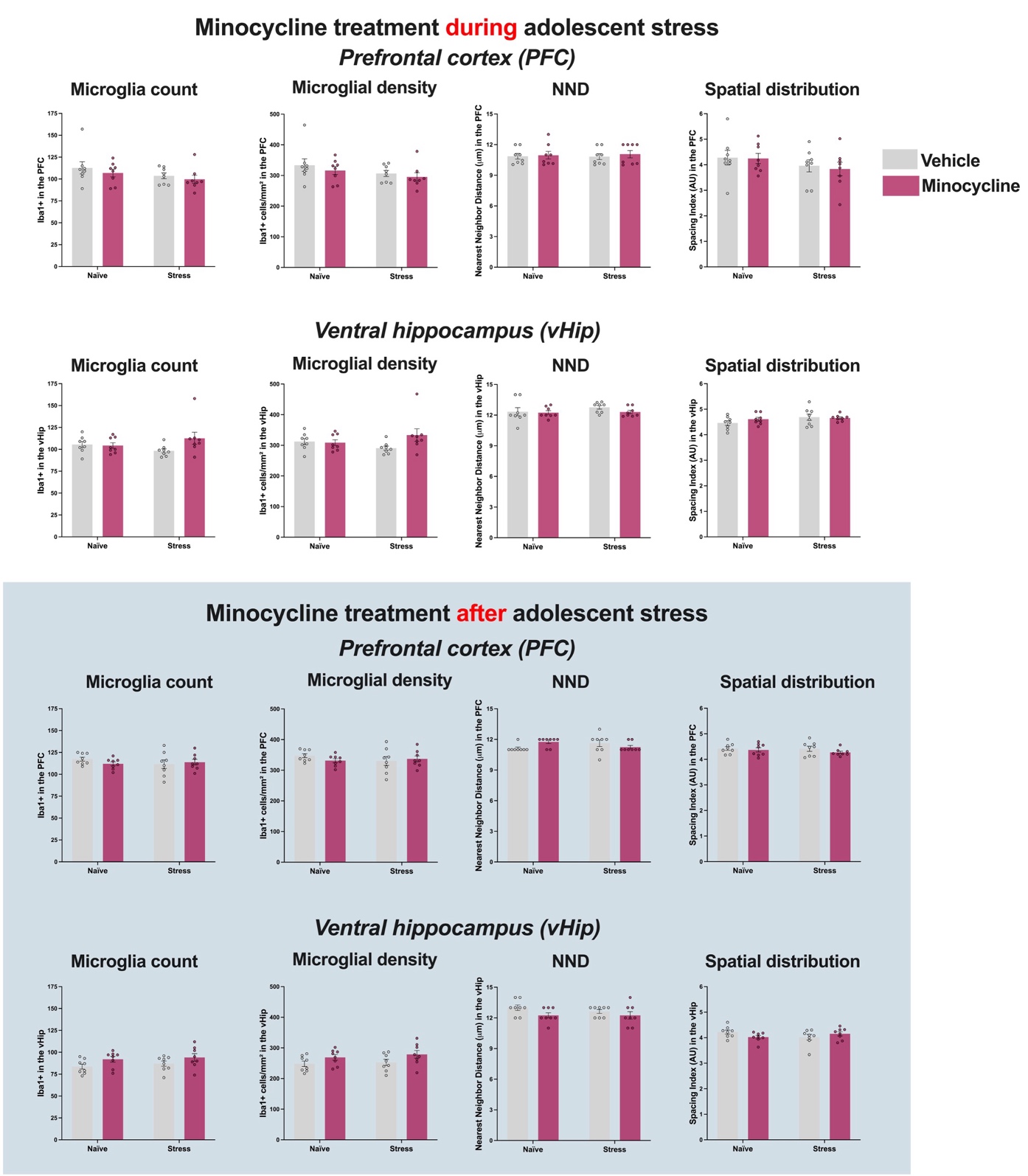

**Supplementary figure 1 -** Adolescent stress and minocycline treatment, regardless of timing (during or after stress), did not alter the number of Iba1+ cells (microglia count) in the PFC and vHip (n=8/group). Similarly, no significant differences were found in **microglial density**, nearest neighbor distance (NND)**,** or the **spacing index** in either brain region.
